# Digital Light Processing 3D printing for biological applications of polydimethylsiloxane-based microfluidics

**DOI:** 10.1101/2022.09.28.509779

**Authors:** Matthew D. Poskus, Tuo Wang, Yuxuan Deng, Sydney Borcherding, Jake Atkinson, Ioannis K. Zervantonakis

## Abstract

Soft lithography microfluidics offer many benefits over conventional biological assays; however, the impact this field is inhibited by the lack of widespread adoption of this technology in part due to prohibitive cost and fabrication time. Recent improvements in three-dimensional (3D) printing technologies such as digital light processing (DLP) printing offer a cost-effective and rapid prototyping solution to microfluidic fabrication. Limited information is available about how 3D printing parameters and resin cytocompatibility impact the performance of 3D printed molds for fabrication of polydimethylsiloxane (PDMS)-based microfluidics for cellular studies. Using a low-cost, commercially available DLP 3D printer, we assess the cytocompatibility of several resins, optimize printer settings and characterize minimum feature size of our system. We demonstrate the applications of DLP printing for soft lithography microfluidics by developing four assays to characterize cell viability, drug response, establish concentration gradients, and monitor live-cell 3D invasion into a hydrogel.

## Introduction

Microfluidics are characterized by their sub-millimeter (<1000*μ*m) features and fluidic channels^1^. Due to their low cost, high sensitivity, and small reagent volumes^2^, and control of cellular environment^3^, microfluidics are widely used in biological applications such cell migration studies^4^, drug sensitivity^5^, and modeling cellular response to fluid flow^6^. Despite these benefits, microfluidics has yet to be widely adopted for biomedical research, in part due to higher cost^7^ and fabrication barriers of entry that may make it impractical for researchers to easily implement^8^. Standard microfluidic mold fabrication using SU-8 mold lithography can be time-consuming (hours to days)^3,9^ and requires specialized training and facilities (cleanroom) to fabricate molds^10,11^.

Additive manufacturing is a promising recent technology in the biomedical field that has several advantages over standard microfluidic fabrication methods^12–15^. Specifically, 3D printing has lesser cost and fabrication time compared to SU-8 mold lithography^16,17^ and does not require a cleanroom^18^. These advantages make 3D printing suitable for rapid prototyping that may accelerate design development^19,20^. 3D printing offers greater design flexibility compared to SU-8 mold lithography by permitting fabrication of truly 3D structures rather than the planar geometries typical of photolithography^21,22^ and allows users to either directly print microfluidics or generate molds for soft lithography^18^. Digital light processing (DLP) is a promising 3D printing technology that offers greater resolution (18-250*μ*m resolution)^23,24^ and improved surface quality compared to fused deposition modeling 3D printing methods^15,17,25^. Briefly, this method uses light generated either from a projector or LCD screen to project a 2D image onto a photopolymerizing resin, locally curing a layer of resin in these illuminated regions. The part is formed layer-by-layer on a build stage until the completed 3D structure is formed^9,26^. Because entire layers of the part can be formed at the same time, DLP is faster than other 3D printing technologies^26^.

Current challenges with this technology include optimizing printer resolution and resin toxicity, as uncured resin components may be cytotoxic^27–29^. Commercial printer resolution is a limiting factor for many researchers^30^. While advancements have been made in analyzing the physical limitations and biocompatibility of resins^25,26,31,32^, few studies have characterized the performance of polydimethylsiloxane (PDMS) microfluidics, a ubiquitous material used in biomedical applications^33^, fabricated via soft lithography using 3D printed molds. Our work addresses these challenges by systematically characterizing the impact of printing protocol and resin on PDMS microfluidic feature resolution and cell viability, respectively. Herein, we demonstrated the impact of universally relevant DLP printing parameters (layer height, part orientation, and exposure time) to optimize printer settings and identify feature resolution limits for multiple geometries. We utilized three relevant microfluidic geometries: microwells, gradient generators, and hydrogel-forming devices to demonstrate the capabilities of a low-cost, commercially available 3D printer in cell viability, drug response and 3D cell migration assays.

## Methods

### 3D Printed Mold Fabrication

Computer-aided design (CAD) models for all molds are generated using Autodesk Inventor (Autodesk, USA) and directly imported into the 3D printer slicer program Chitubox (Chitubox, China) for printing preparation. Resin-specific default printing profiles were used unless otherwise specified. “Medium” support settings were used and supports were automatically generated using the “+All” setting. Components were printed using a Phrozen Sonic Mini 4K (Phrozen Technology, Taiwan) with screen protector (BulletBrandCompany, USA). Z-Calibration of the printer was performed per manufacturer instructions prior to each print. Resin (Phrozen Technology, Taiwan) was filled to approximately 1/3 the height of the resin vat. After printing is completed, excess resin is removed from the components using compressed air. Unused resin is filtered using 150*μ*m paper strainer (Shanqian, China) to be reused. Components are placed in an ANYCUBIC Wash and Cure station (Anycubic, China) to wash for 10 minutes. After washing, parts are removed from the printing stage using a metal spatula and placed in a plastic bag filled with 70% isopropanol/30% deionized (DI) water. The bag is placed in an ultrasonic cleaner (Kaimashi, China) for sonication in a water bath for five minutes. Parts are subsequently dried with compressed air before placement in the ANYCUBIC Wash and Cure Station to cure for 60 minutes. Excess resin is removed from the building plate and the build area is resurfaced by sanding using 60-grit sandpaper for 10-20 seconds prior to printing again. After curing, the parts are placed in an oven (Hybaid, USA) for 48 hours before returning to storage at room temperature.

### PDMS Fabrication

Sylgard 184 PDMS is vigorously mixed in a 10:1 ratio elastomer base to curing agent by mass and placed in a vacuum desiccator for 1 hour to remove bubbles. PDMS is poured into molds and placed overnight in an oven (Hybaid, USA) overnight to cure before use. PDMS was removed from molds using a hobby knife. Debris is removed from devices prior to use with tape (Scotch, USA). The first pour of PDMS from all molds is discarded to account for any potential uncured resin transferring to PDMS.

### Cell Culture

Breast cancer cell lines BT474 and MDA-MB-231 were generously provided by Dr. Dennis Slamon, University of California Los Angeles, Los Angeles, CA,. BT474 and MDA-MB-231 cell lines were engineered to express H2B-mCherry by retrovirus and subsequently sorted using fluorescence-activated cell sorting (FACS) to select for mCherry+ cells. Cells were grown in Roswell Park Memorial Institute (RPMI) 1640 media (Corning, USA) supplemented with 10% heat-inactivated fetal bovine serum (HI-FBS) (Avantador, USA) and 1% penicillin (100 units/mL)/streptomycin (100*μ*g/mL). (Gibco, USA). Cells were cultured in a humidified incubator at 5% CO_2_ and 37°C.

### Microwell Viability Assay

3D printed molds were fabricated using 50*μ*m layer height and 55° orientation settings. PDMS was sterilized by autoclave (30 minutes at 121°C wet cycle followed by 30 minutes at 121°C gravity cycle). All molds were plasma treated using a plasma cleaner (Harrick Plasma, USA) prior to use to enhance surface hydrophilicity and prevent bubble formation in microwells. A reservoir for SU-8 microwells was created by plasma binding a second layer of PDMS to the surface of the microwells. Photolithography was used to produce micropatterned SU-8 molds (MicroChem, USA). Cells were collected by trypsinization (0.05% Trypsin, Corning, USA) and seeded at 33,000 cells/cm^2^ in 1.75mL for 3D printed microwells, 500*μ*L in SU-8 microwells, and 200*μ*L in black 96-well plates (Greiner Bio-One, Germany). To monitor cell death cells are cultured in media containing 100nM Sytox Green (Invitrogen, USA). 100nM Paclitaxel (Selleck Chemical LLC, USA) added upon cell seeding in treatment conditions. Media is replenished daily as needed. Imaging was performed using a Nikon Ti2 microscope (Nikon, Japan) equipped with a live-cell imaging stage (Tokai Hit, Japan). Analysis of fluorescence images was performed using an Ilastik^34^ machine learning pipeline and CellProfiler^35^ to classify individual cells as alive or dead. Viability was calculated as the fraction of alive cells of total cells (alive and dead). Replicates represent individual biological replicates. Error bars represent standard deviation.

### Printing Parameter Optimization and Feature Size Characterization

3D printed molds were fabricated using 50*μ*m layer height and 55° orientation settings unless otherwise specified. PDMS poured from molds was imaged via widefield microscopy using a Nikon Ti-2 (Nikon, USA). Microwell aspect ratio was computed as the ratio of height to width using ImageJ. Replicates represent randomly sampled microwells from one PDMS microwell array. For feature size characterization, 3D printed molds were fabricated using 10*μ*m layer height and 29° XY orientation settings. SU-8 wafers were fabricated in-house (University of Pittsburgh, USA). PDMS cast from molds was plasma treated then bonded to a glass slide for imaging via plasma treatment (Harrick Plasma, USA). PDMS from molds was imaged using a Nikon Ti2 (Nikon, Japan) microscope. ImageJ was used to quantify the length of each side of the features. The mean side length for all edges is compared to the nominal feature side from the corresponding CAD model. Error bars represent standard deviation.

### Gradient Generator Assay

Inlet and outlet ports in the device are punched using a 2mm biopsy punch (Miltex, USA). Devices are bound to 24×60mm #1 coverglass (VWR, USA) via plasma treatment using a plasma cleaner (Harrick Plasma, USA). A solution of DI water or DI water containing 4*μ*g/mL 10kDa AlexaFluor-647-conjugated dextran (Invitrogen, USA) and 1:20,000 dilution of red fluorescent 1*μ*m carboxylate-modified microspheres (Invitrogen, USA) was prepared. These solutions were aspirated into 10mL syringes (BD, Switzerland) connected to microbore tubing (Masterflex, USA) that were connected to the microfluidic device via luer connectors (Qosina, USA). The outlet port was connected to tubing that fed into a waste beaker. Tubing was directly inserted into inlet/outlet ports to form a leakproof connection. Syringe pump-driven flow was established using a two-channel syringe pump (Harvard Apparatus, USA) at the prescribed flow rate. Flow was established for seconds to minutes depending on the flow rate to achieve steady state for each condition prior to imaging. Imaging was performed on a Nikon Ti2 (Nikon, Japan) microscope. Channel intensity was quantified using NIS-Elements (Nikon, Japan) and ImageJ measuring the mean intensity in each channel.

### 3D Invasion Assay

Ports in the device are punched using a biopsy punch (Miltex, USA). Devices are sterilized via autoclave (30 minutes at 121°C gravity cycle). Device are attached to 22×40mm #1 coverglass (VWR, USA) via plasma treatment using a plasma cleaner (Harrick Plasma, USA). Once bound, devices are incubated at 80°C for 48 hours to return the PDMS surfaces to a hydrophobic state. The central channel of devices is filled with buffered collagen type I (Corning, USA) at a concentration of 1mg/mL and incubated for 30 minutes at 37°C to polymerize. MDA-MB-231 breast cancer cells are subsequently seeded at a concentration of 0.5M cells/mL into the device (40*μ*L) in either 0% FBS media (starvation) or 10% FBS media (full media). Devices are placed on an incubator stage (Tokai Hit, Japan) for live-cell imaging using a Nikon Ti2 microscope (Nikon, Japan). Cells were imaged every 15 minutes for 8 hours. Devices were fixed using paraformaldehyde (Electron Microscopy Sciences, USA) after 24 hours. Cell migration and invasion analysis was performed using ImageJ TrackMate^36^. Confocal imaging was performed using a Zeiss LSM700 (Carl Zeiss, Germany) with 5*μ*m z-step. Error bars represent standard deviation.

## Results

### Fabrication of printed molds using DLP 3D printing

A commercially-available Phrozen Sonic Mini 4K DLP printer (**Figure 1A**) was used to fabricate 3D printed molds. Briefly, CAD models (i.e., STEP files) are imported into Chitubox slicer software for conversion to 3D printer format and to set 3D printing parameters (e.g., layer height, exposure time, supports, etc.). Printing is performed by loading the slicer file onto the printer via USB drive and filling resin vat. The ultraviolet (UV) liquid crystal display (LCD) screen below the resin vat illuminates specific pixels to project an image of the slice onto the resin to induce local photopolymerization of the resin to form one cured layer. This process is repeated for each layer. After curing, the printed parts undergo a series of post-processing steps including two washing steps, a UV postcure, and thermal postcure to prepare the mold for soft lithography and to limit potential PDMS curing inhibition (**Figure 1B**). PDMS is poured into the molds to produce microfluidic devices (**Figure 1C**).

**Figure 1:**
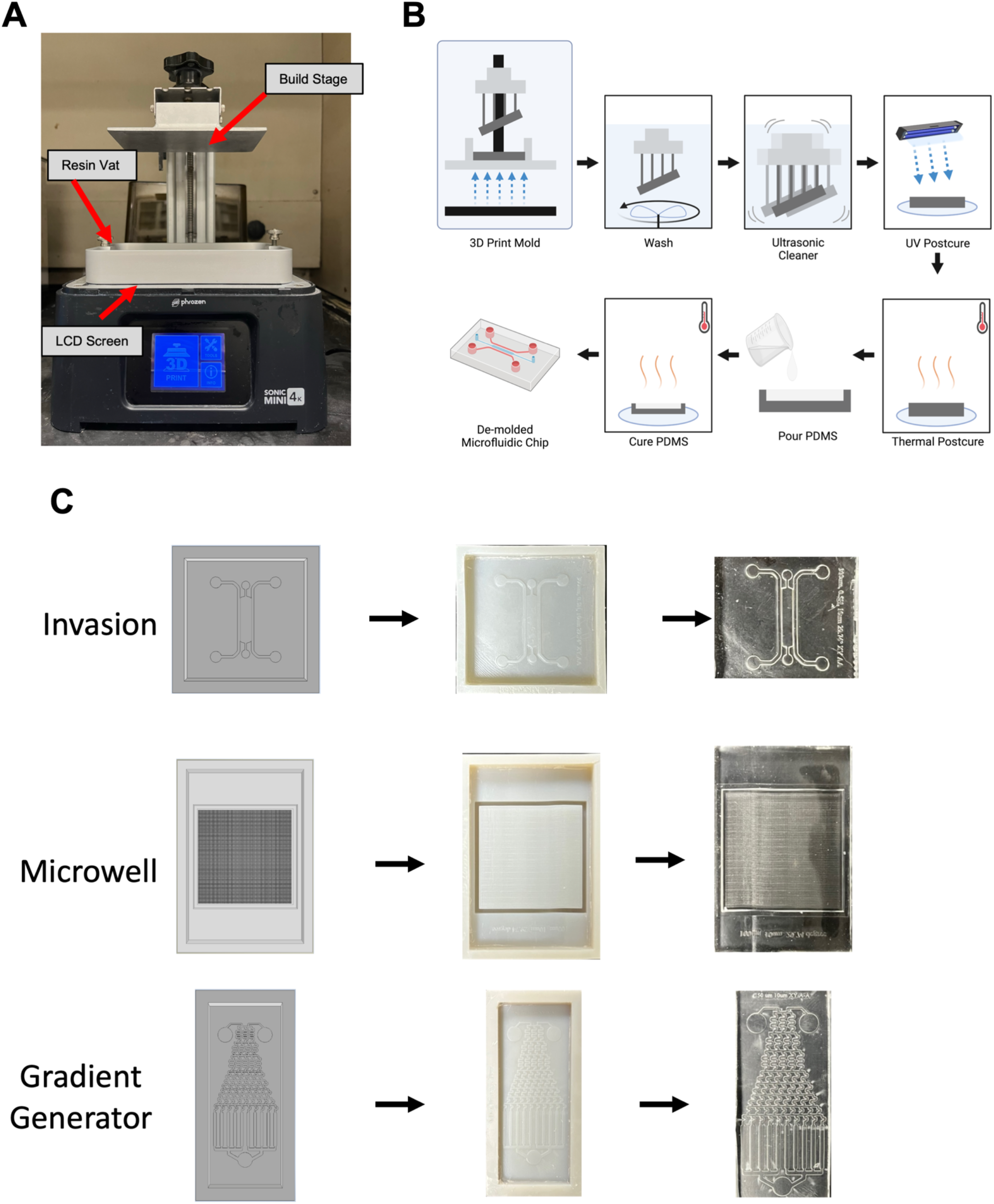
Process overview for DLP 3D printing of microfluidic devices. (A) Commercially available Phrozen Sonic Mini 4K printer used for mold for fabrication. (B) Workflow for fabrication of microfluidic devices using DLP 3D Printing. (C) Representative images of CAD model (left), 3D printed mold (middle), and microfluidic chip (right) for geometries tested.

### Characterization and biocompatibility of DLP resins

We identified several commercially available resins that may be suitable for 3D printing molds for soft lithography. We limited our selection to resins recommended for our printer, as we reasoned these to have the greatest performance without exhaustive testing and validation. We selected five resins with a range of mechanical properties for our studies (**Table 1**). To assess the performance of these resins, we first compared the quality of an array of 100×100*μ*m microwells fabricated from molds of each resin material (**Figure 2A**). Each resin was able to produce distinct microwell features (**Figure 2B**). We noted the beige dental resin developed cracks in the mold after the thermal postcure stage (**Supplementary Figure 1A-B**). We next seeded BT474 tumor cells into 3D printed microwells to assess biocompatibility of PDMS from each of the resins for up to four days. Viability of tumor cells in microwells exceeded 85% viability for all resins (**Figure 2C-D**). To ensure cell response to therapy in 3D printer PDMS mimics other *in vitro* models, we treated BT474 tumor cells with 100nM paclitaxel for 4 days to confirm cellular response to drug treatment response in microplates (**Figure 2E-F**). Indeed, drug response was similar between the standard microplate and our 3D printed microwell assay, as measured up to four days after treatment.

**Table 1:** Comparison of mechanical and thermal properties of tested resins.

| Resin | Viscosity (cps) | Ultimate Tensile Strength (Mpa) | Tensile Modulus | Heat Deflection Temperature (C) |
| --- | --- | --- | --- | --- |
| Nylon Green | 850-905 | 24 | 600 | Not Specified |
| TR250LV | 180-280 | 25 | 900 | 100-120 |
| Rock Black | 70-170 | 30 | 419 | 97 |
| Rapid Black | 70-170 | 15 | 110 | Not Specified |
| Beige Dental | 700-800 | Not specified | Not specified | Not specified |

**Figure 2:**
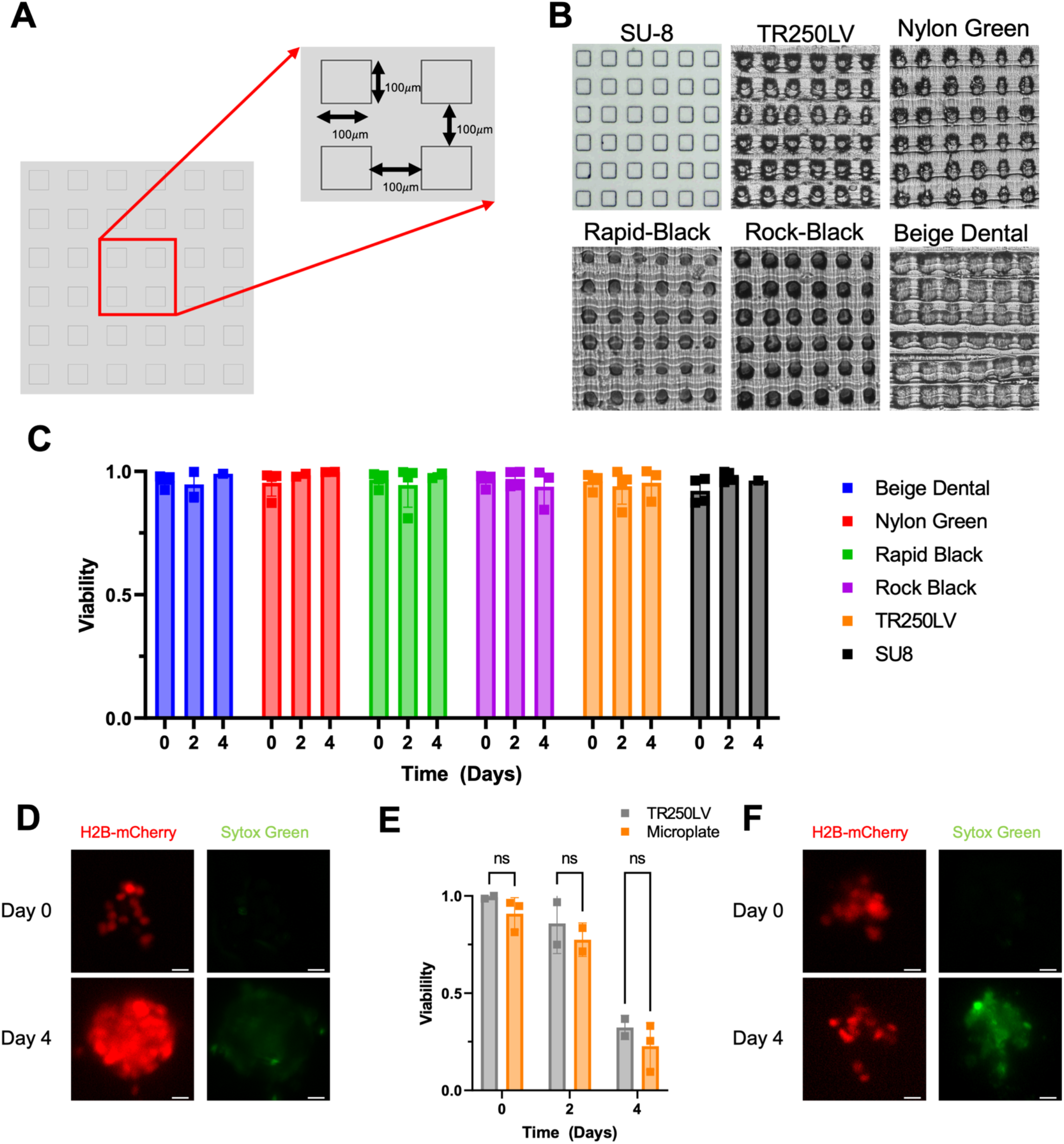
Cell viability and drug response of tumor cells in microwells fabricated using DLP 3D printing. (A) Annotated CAD model of microwell array dimensions. (B) Representative images of PDMS from 3D printed molds for tested resins. (C) Viability after 4 days of cancer cells seeded in PDMS from 3D printed molds. (D) Representative images of tumor cells seeded in microwells after 4 days. (E) Viability after 4 days of cancer cells seeded in PDMS of treatment with 100nM paclitaxel. (F) Representative images of tumor cells seeded in microwells after paclitaxel treatment. Scale bars represent 25*μ*m.

### Printing Parameter Optimization

To systematically characterize the impact of each printing parameter on printing quality, the TR250LV resin was selected owing to its heat deflection temperature and mechanical properties (**Table 1**). As biocompatibility of PDMS from all resins seemed comparable, we reasoned that these properties may increase mold longevity after repeated use and heat-cycling when curing PDMS. We first explored the impact of layer height—the thickness of an individual layer of the mold—on PDMS quality by examining microwells of PDMS cast from molds printed at 50*μ*m and 10*μ*m. Qualitatively, the PDMS from molds printed at 10*μ*m had fewer and less severe aliasing lines (**Supplementary Figure 2A**) caused by adjacent printed layers compared to molds printed at 50*μ*m. Furthermore, the aspect ratio, measured as the ratio of length to height of individual microwells, was significantly closer to the nominal value of unity for the designed microwells (**Supplementary Figure 2B**). Therefore, we continued all future prints with 10*μ*m layer height. The impact of print orientation was next explored. Molds were printed at 55°, 29°, or 15°, which represent the angle of a “staircase” of voxels in which the length of the step represents the (fixed) pixel size (35*μ*m) and the height of the step represents layer height at 50, 20, and 10*μ*m. We reasoned that this angle would minimize aliasing for the given pixel size and layer height. PDMS cast from molds printed at these angles was examined (**Supplementary Figure 2C**). At the 29° and 15° orientation, aliasing is reduced. However, at shallower printing angles the microwells become elongated rather than square and have an increased aspect ratio. Since both the 29° and 15° microwells have minimal aliasing, the 29° printing angle was considered superior due to its lesser elongation (**Supplementary Figure 2D**). However, PDMS from all molds was not flat due to deformation of the mold. This was corrected by orienting molds along two axes (about the x- and y-axis) rather than one (about the x-axis) (**Supplementary Figure 2E**). This resulted in increased distortion of microwells (**Supplementary Figure 2F**); however, these microwells had lesser elongation (**Supplementary Figure 2G**). Finally, the impact of layer exposure time on PDMS cast from molds was examined. At a reduced exposure time, microwells are undeveloped (**Supplementary Figure 2H**). These analyses identified optimal printing settings of: 10*μ*m layer height, 29° orientation along two axes, and default exposure time (2.5s) for TR250LV resin. The final printing parameters selected for all prints were: 10*μ*m layer height, 29° two-axis printing orientation, and 2.5 second exposure time.

### Feature Size Characterization

We next asked what is the minimum feature size that can be resolved by our printer. To this end, we printed positive and negative features of varying shape and size and compared the measured and nominal feature sizes. A series of lines of decreasing thickness were printed and measured using ImageJ (**Figure 3A**). The measured feature size closely matched the nominal feature size (**Figure 3B**). All lines down to 100*μ*m were resolved in the PDMS. Hexagonal embossed and debossed features were printed and the PDMS feature size was compared to nominal feature size (**Figure 3C-D**) ranging from 35-600*μ*m in the length of each side. While all features were resolved in the SU-8 PDMS, only hexagons with lengths down to 150*μ*m were sufficiently resolved to measure.

**Figure 3:**
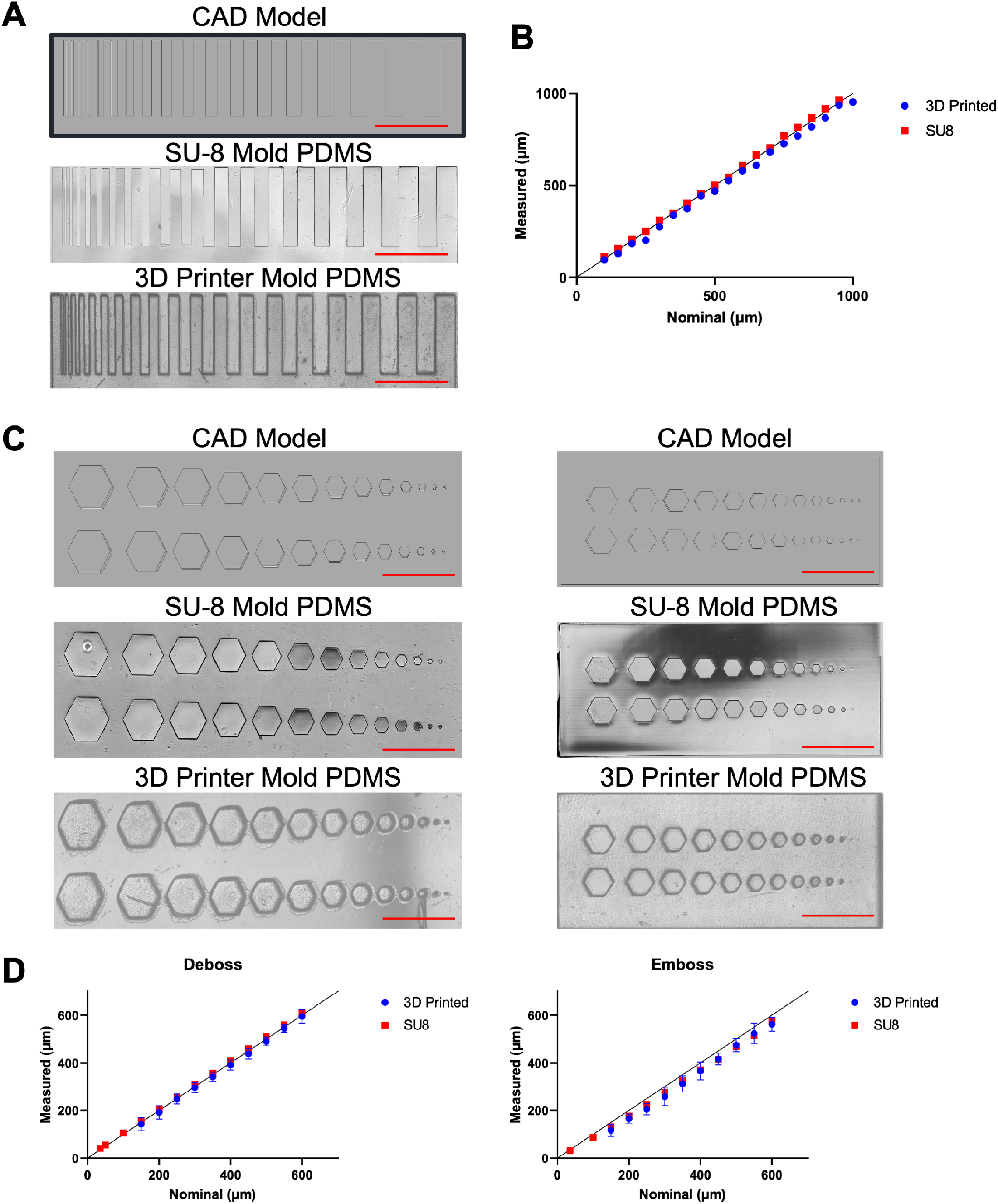
Minimum feature size of DLP 3D Printed microfluidics. (A) Representative images of lines of varying width. (B) Quantification of nominal vs. measured feature size of PDMS from SU-8 and 3D printer molds (C) Representative images of debossed (left) and embossed (right) hexagons. (D) Quantification of feature size for debossed (left) and embossed (right) hexagons. Scale bars represent 1mm.

### Microfluidic Gradient Generators

We developed a microfluidic concentration gradient generator to demonstrate the capacity of our system to fabricate microfluidic channels and sustain flow. The microfluidic concentration gradient generator we tested consists of two inlets, one outlet and ten channels across which a gradient is formed (**Figure 4A**). A solution of fluorescently labeled (AlexaFluor 647) dextran and red fluorescent beads suspended in DI water was perfused through top inlet and pure DI water in the other inlet to form a gradient. A two-channel programmable syringe pump was used to ensure identical flow rates into each inlet to balance flow. Flow was established at 0.4-50 *μ*L/min and images of the device were acquired using fluorescence microscopy. At low flow rates, a gradual change in dextran concentration occurred across the device channels, whereas at high flow rates a step-change in concentration occurred (**Figure 4B-C**). Fluorescent beads were used to probe particle streaklines within the device to ensure channels did not leak. At both low and high flow rates, particle streaklines did not deviate from the channel, suggesting the channels maintain strong adhesion to glass even at high flow rates (**Figure 4D**).

**Figure 4:**
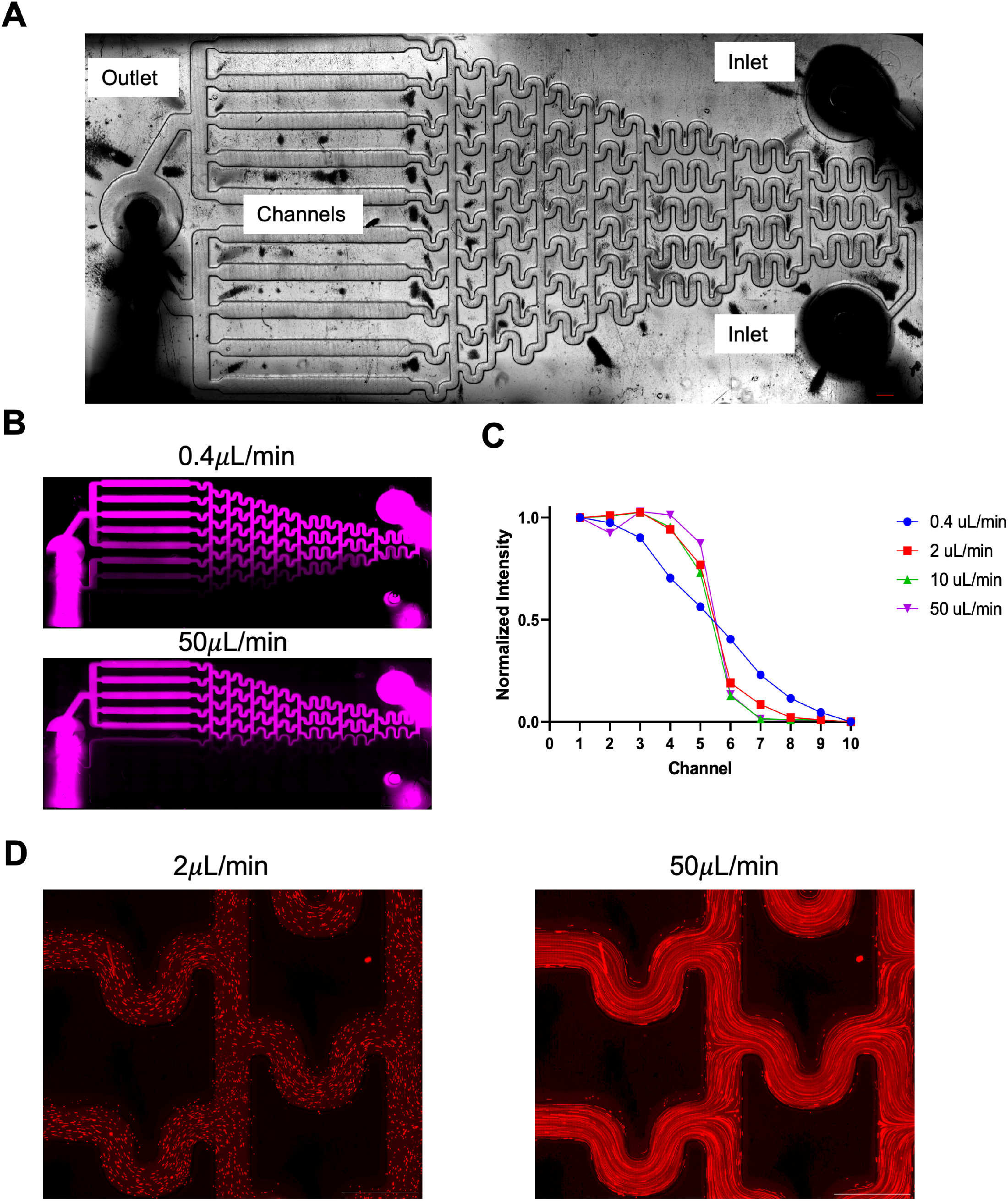
Microfluidic gradient generator fabricated using DLP 3D printing. (A) Schematic of microfluidic gradient generator. (B) Dextran gradient produced by perfusing at 0.4*μ*L/min (top) and 50*μ*L/min (bottom). (C) Intensity of fluorescent dextran measured in each channel for tested flow rates. (D) Fluorescent beads flowing through gradient generator at 2*μ*L/min (left) and 50*μ*L/min (right). Scale bars represent 1mm.

### 3D Cancer Cell Invasion in a Collagen Gel

A microfluidic cancer cell invasion assay was performed to assess device performance in sustaining embedded hydrogel and tracking single-cell migration. Briefly, collagen was polymerized in the center microfluidic channel and MDA-MB-231 breast cancer cells were seeded in the outer channel in either starvation or full media conditions (**Figure 5A**). Cancer cells were imaged every 15 minutes to track individual cells for up to 8 hours. After the first 8 hours, cancer cells invaded into the gel in full media condition but not starvation (**Figure 5B**). We tracked the migration trajectory of individual cells and asked whether the migration speed varied between full media and starvation conditions. For 0-4- and 4-8-hour windows, the migration speed of cells in full serum was greater than cells in starvation conditions (**Figure 5C**). Additionally, the total displacement of cells in the full starvation was also greater for full serum conditions (**Figure 5D**). We also confirmed the distribution of invaded cells in the 3D collagen hydrogel via confocal imaging (**Figure 5E**).

**Figure 5:**
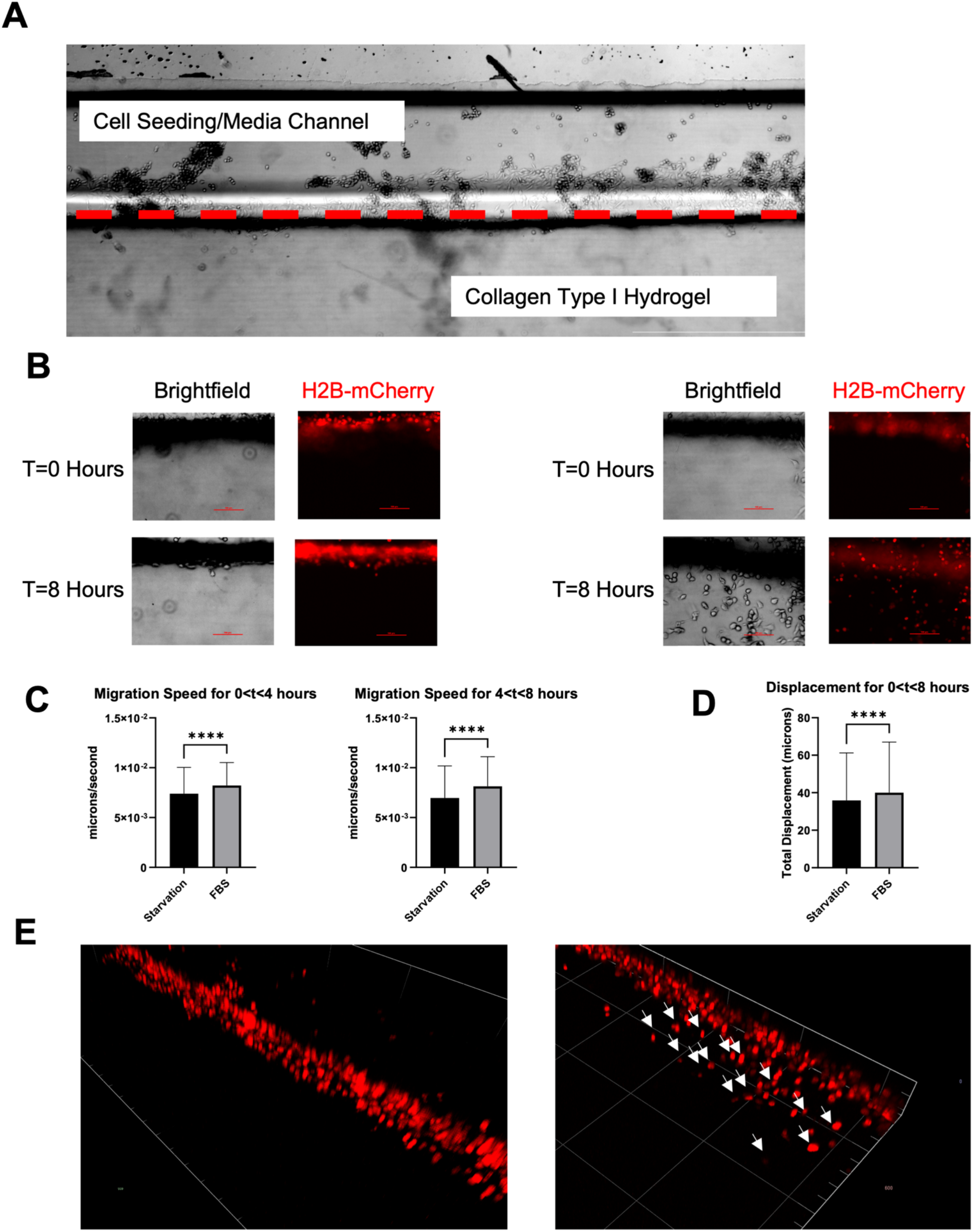
Cancer cell invasion invasion into collagen type I. (A) Schematic of invasion assay. Red dashed line indicates hydrogel invasion interface. (B) Representative images of MDA-MB-231 cancer cells stimulated with starvation (left) or full media (right) after 8 hours. (C) Quantification of migration speed of single cells between 0-8 hours. (D) Quantification of cancer cell displacement after 8 hours. (E) Confocal imaging of invading cancer cells in starvation (left) or full media (right) conditions. White arrows indicate examples of invading cells.

## Discussion

3D printing is a relatively recent technology that has gained enormous attention in the biomedical field^15,37^ due to its low cost, capacity for truly 3D features, and potential for rapid prototyping^9^. However, this technology is still in its infancy^38,39^. In addition to current technical limitations that must be overcome (e.g., improved printer resolution^30^), biocompatibility of these 3D printed materials is an unceasing consideration for cellular studies that remains to be rigorously explored. Although several recent studies have characterized the performance of 3D printed microfluidics, it is not currently clear how these results translate to microfluidics produced using soft lithography from 3D printed molds. To fulfill this need, herein we utilized a low-cost (less than $300 USD), commercially available DLP resin printer to produce microfluidics using soft lithography. This printer was selected due to its affordability compared to other 3D printing systems which can range from $2,000-$10,000 USD^25,40^, which may be prohibitively expensive for some researchers. We selected several commercially available printer resins and characterized the quality of PDMS microwells cast in molds of each resin. Notably, all tested resins produced well-defined 100×100*μ*m microwells which is superior in resolution compared to previous reports of minimum feature size using commercially available printers^19^. All tested resins demonstrated sufficient (<85% viability) when used for soft lithography and is consistent with other reports^41^. These results are a promising first-step of understanding biocompatibility of 3D printed molds for soft lithography microfluidics. Owing to the sensitivity of 3D printer resolution to environmental, mechanical, and optical factors^26,42–44^, we empirically tested various printer settings to optimize feature quality. While our optimal printing configuration yielded slight distortion of the microwell features, we did not note any significant distortion in subsequent feature size characterization studies nor in our applications of gradient generators and cell invasion. Therefore, we attribute this distortion to the relative scale of the microwells (100×100*μ*m) compared to our printer pixel size (35*μ*m); the microwell features may not be properly resolved when discretized into relatively large voxels (35×35×10*μ*m) printed on an angle. Despite these challenges, we report that our printer can resolve straight lines down to 100*μ*m and 150*μ*m hexagonal features, which is an improvement from resolution offered by other commercially available systems^45,46^. Others have reported impressive resolutions as small as 20*μ*m through customized resin formulations and equipment^47–51^ which highlights the potential of 3D printing for truly microscale features. However, these resources are not widely available to researchers. Recent advancements in commercially available printers that achieve resolutions of 22*μ*m are bridging the gap in resolution to offer affordable access to high-resolution printing^52^.

## Acknowledgements

The authors acknowledge grant support from the US National Institutes of Health (R00 CA222554 to I.K.Z. and T32 EB001026 to M.D.P.). Some figures created with Biorender.com.

---

**Supplementary Figure 1:**
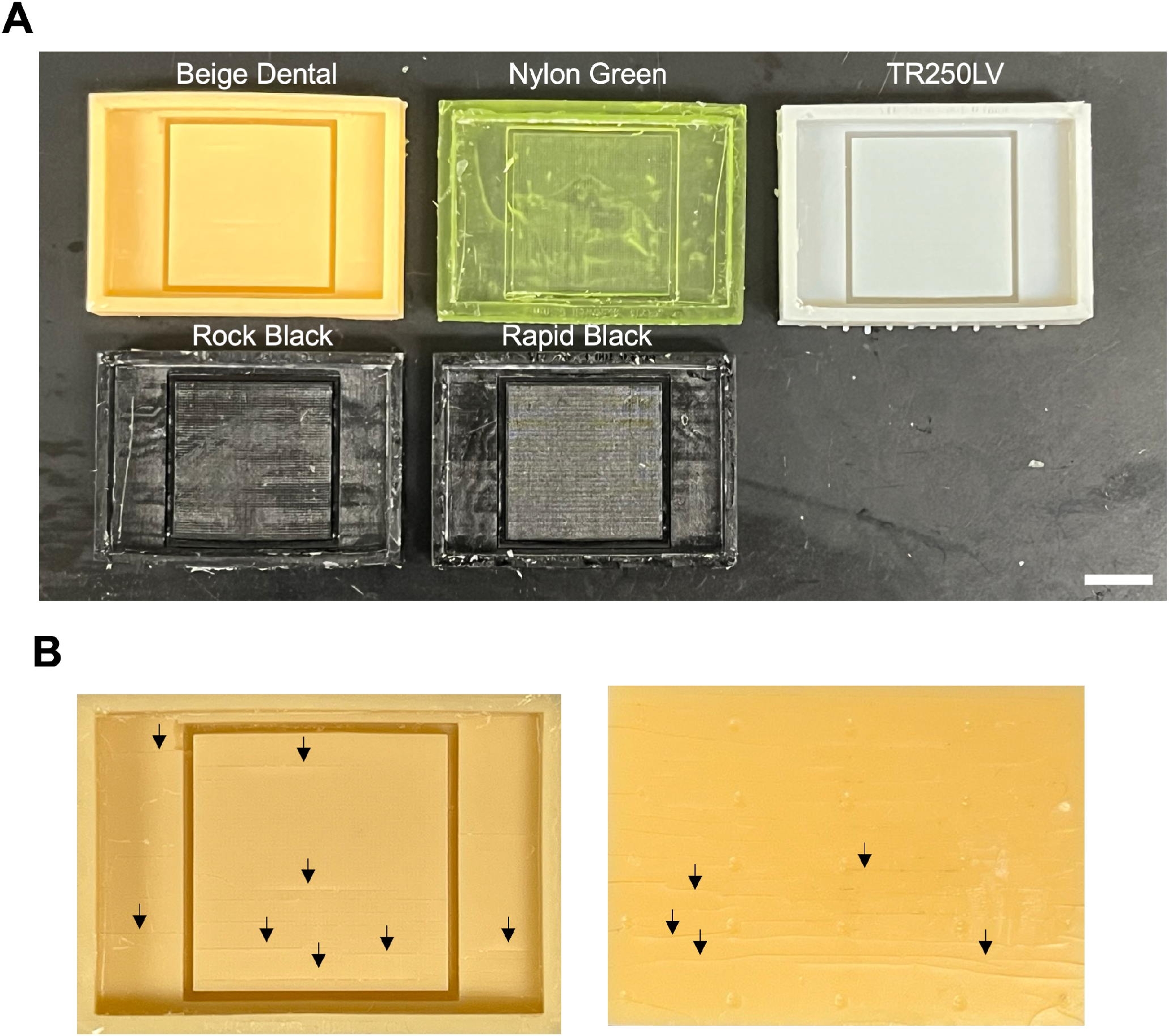
Microwell molds after thermal postcure stage. (A) Images of DLP 3D printed molds of five tested resins. Scale bar represents 1cm. (B) Cracks formed in beige dental resin on the top (left) and back (right) of mold after thermal postcure. Black arrows indicate examples of cracks.

**Supplementary Figure 2:**
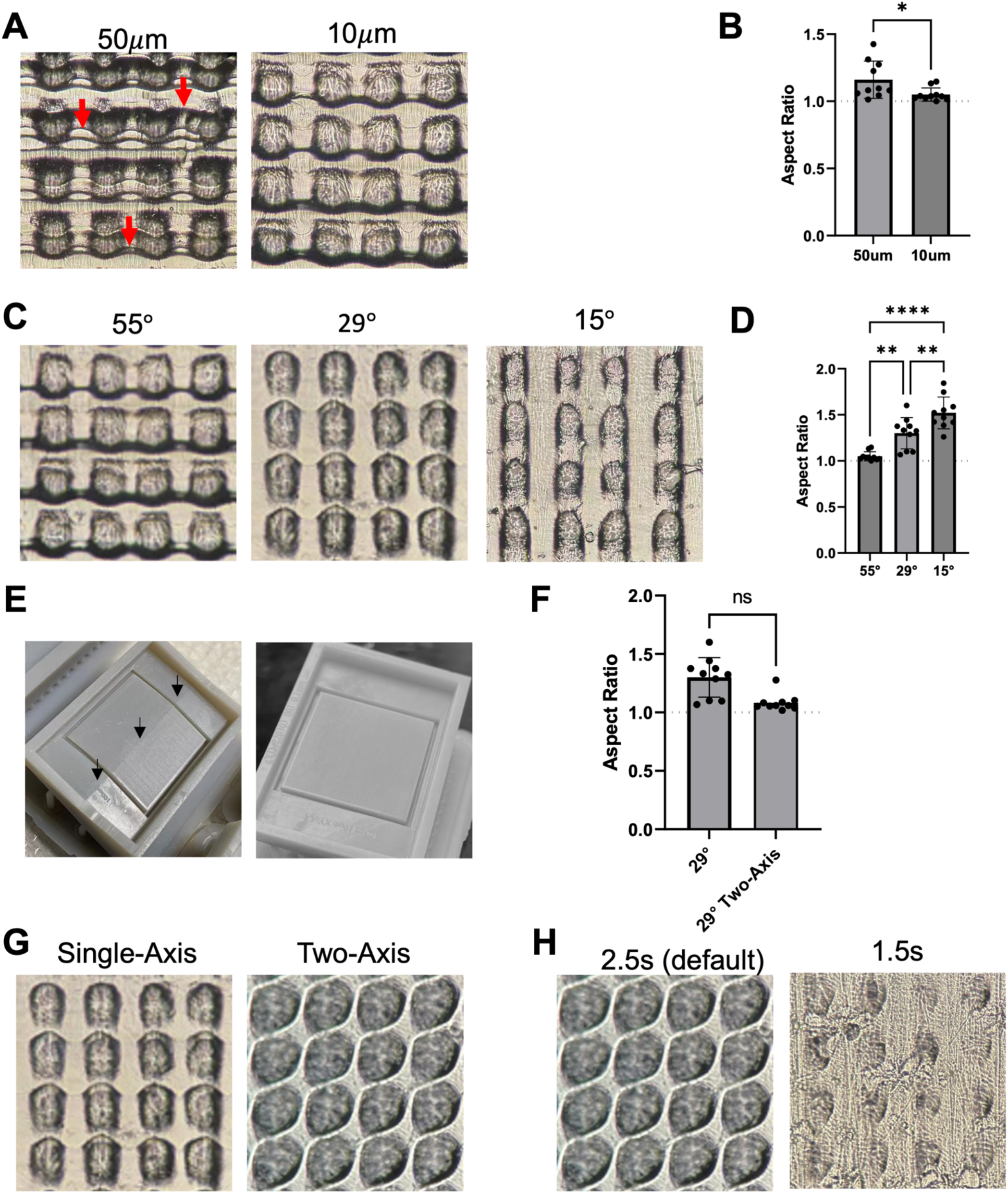
Optimization of DLP printing parameters to maximize feature quality. (A) Comparison of PDMS molds at 50*μ*m (left) or 10*μ*m (right) layer height. Red arrows indicate examples of horizontal aliasing lines. (B) Quantification of PDMS microwell aspect ratio between 50*μ*m and 10*μ*m molds. (C) Comparison of PDMS from molds printed at various orientations. (D) Quantification of aspect ratio for various printing orientations. (E) Mold deformation during single-axis (left) is corrected by two-axis orientation (right). (F) Comparison of PDMS from molds printed at either single-axis (left) or two-axis orientation (right). (G) Quantification of PDMS microwell aspect ratio between single- and two-axis orientation.

